# What colour are Actinomycetes? We have a tool for that! PhylaChrome - A systematic and deterministic tool for assigning colours that capture taxonomic relationships in microbiome datasets

**DOI:** 10.64898/2026.07.29.741550

**Authors:** Rachel L. Theriault, John Parkinson

## Abstract

Visual representations of data generated from complex microbial communities (i.e. microbiomes) can provide intuitive insights into their structure, function, and dynamics. However, current approaches typically rely on *ad hoc* selection of colour palettes that fail to capture taxonomic relationships and limit cross-study comparisons. PhylaChrome has been developed to systematically and deterministically assign colours to enhance visual separation of taxa, capture taxonomic relationships and facilitate intuitive comparisons across datasets.

## MAIN TEXT

Complex microbial communities (microbiomes) feature thousands of interactions supporting community structure and function. Defining their composition and taxonomic relationships are key to understanding community dynamics in the context of the ecosystems they inhabit. Supporting these studies are intuitive visual representations that help capture these relationships ^1^; a major component is the choice of colour to represent taxa ^2-4^. Colour choice plays a vital role in effective scientific communication but is often overlooked ^5^. Traditionally taxa are distinguished through the application of user defined colour palettes to ordered taxa (for example user-defined or on the basis of abundance) that while aesthetically pleasing, vary across studies and are limited in their ability to capture taxonomic relationships (e.g. ^6, 7^). To address these shortcomings, we have developed PhylaChrome, an R package that generates taxonomically informed colours for visual distinction of bacterial taxa. While attempts have been made to define colours based on taxonomic relationships, they are restricted to a limited set of user-defined taxa ^8^. PhylaChrome systematically and deterministically assigns colours across entire taxonomic lineages. In addition to ensuring that taxonomic relationships are captured through colour, the deterministic approach used by PhylaChrome facilitates intuitive comparisons across studies.

PhylaChrome defines a colour for each bacterial phylum that is converted to the (H)ue (S)aturation and (L)ightness (HSL) space; every taxon below a phylum is assigned the same hue, and S and L values differentiate lower-level taxa (**Figure 1**). The program can be run in two modes: global or dataset-specific. Under global mode, a breadth-first approach is used for parsing a taxonomic tree in which every taxon is assigned a colour such that all children of a parent taxon occupy a region of the SL space defined by the parent. Mapping to the SL space is performed by applying a cryptographic hash function to the scientific name of a taxon that derives four digits which are then used to define S and L values. This mapping based on the taxon name ensures taxa are represented by the same colour irrespective of the dataset (i.e. Actinomycetes will always be represented by the same shade of red). As the SL space bounded by parent taxa decrease in size, colours may be challenging to visually distinguish. To account for this, the user can assign custom colours to taxa of interest (which are subsequently propagated to their children) to ensure they can be readily distinguished.

**Figure 1.**
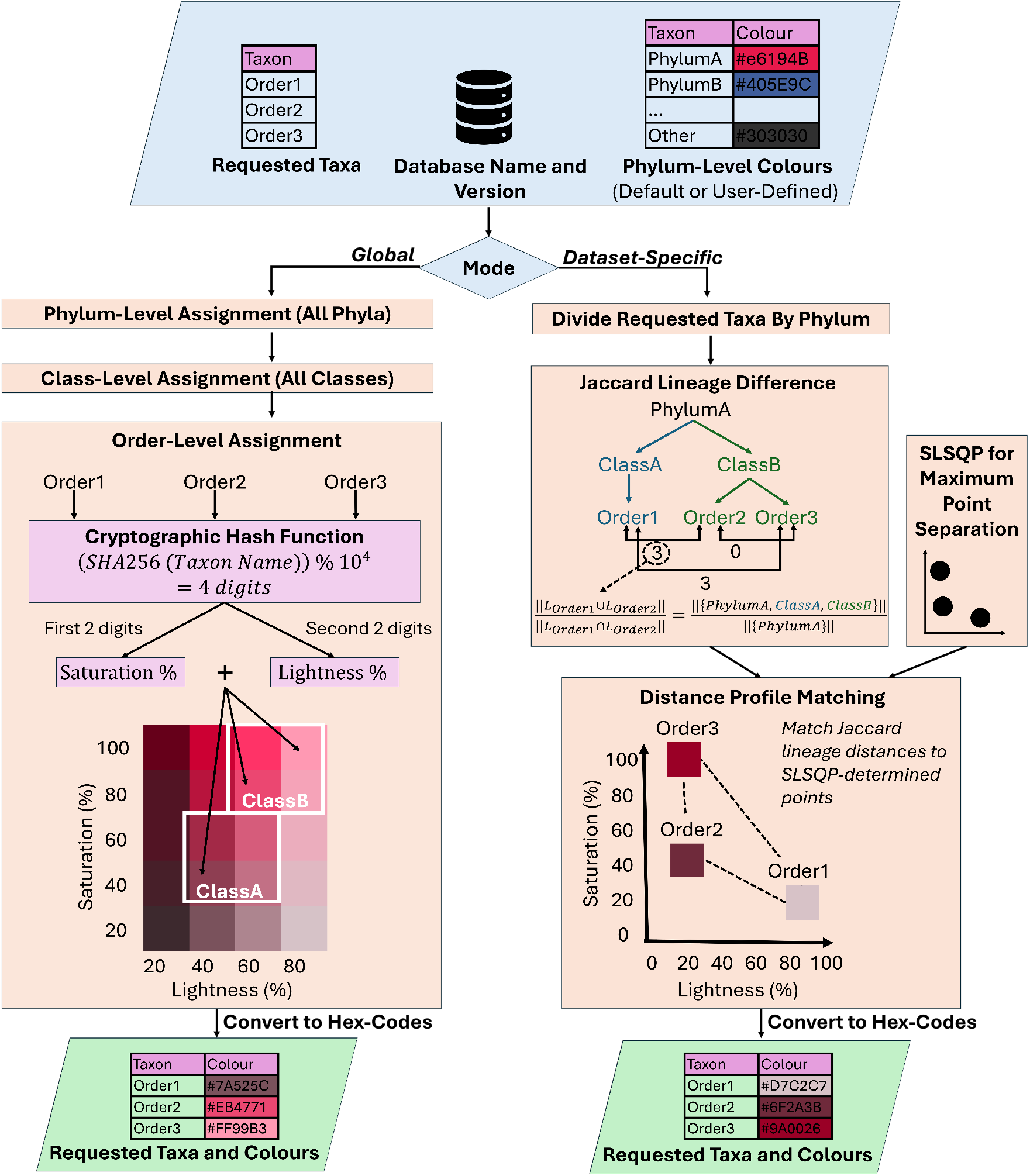
PhylaChrome algorithm for global mode (left) and dataset-specific mode (right) is shown for three requested taxa at the order level. For simplicity, the schematic shows three orders belonging to the same phylum. In practice, requested taxa can include taxa from multiple phyla and/or multiple taxonomic levels.

In dataset-specific mode, as before, all taxa are assigned the same hue as their parent taxon however, present taxa are now assigned SL values that maximize separation within each phylum. For *m* taxa in a phylum, a sequential least squares programming (SLSQP) algorithm is used to generate *m* datapoints that maximize separation in the SL space. The pairwise Jaccard difference between taxon *i* and taxon *j* is calculated using equation (1),

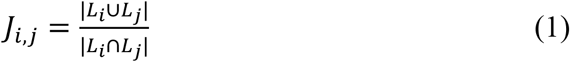

where *L*_*i*_ is the lineage of taxon *i*. A distance profile matching algorithm is then used to map each taxon to a fixed datapoint from SLSQP such that those with similar lineages (based on the number of shared parent nodes) are assigned to points closest in the SL space. Dataset-specific mode produces colours that are more visually differentiable than global mode but only considers present taxa therefore limiting cross-study comparisons.

To demonstrate the utility of PhylaChrome we applied it to nine shotgun metagenomic datasets from different: hosts ^9-11^; human body sites ^12-14^; and natural environments ^15-17^ (**Figure 2; Supplemental Table 1**). After removal of low-quality reads and host sequences (see **Online Methods**), reads were classified using Kraken2 ^18^. Community structure of each dataset was visualized at the order level as a stacked bar chart of relative abundances. Colours were assigned to each taxon for all nine datasets using PhylaChrome global mode. In addition, for three representative datasets we generated charts using PhylaChrome dataset-specific mode as well as the default rainbow colour scheme from R mapped onto taxa ordered by phylum, class, and order membership extracted from the corresponding PhyloSeq^6^ OTU table.

**Figure 2.**
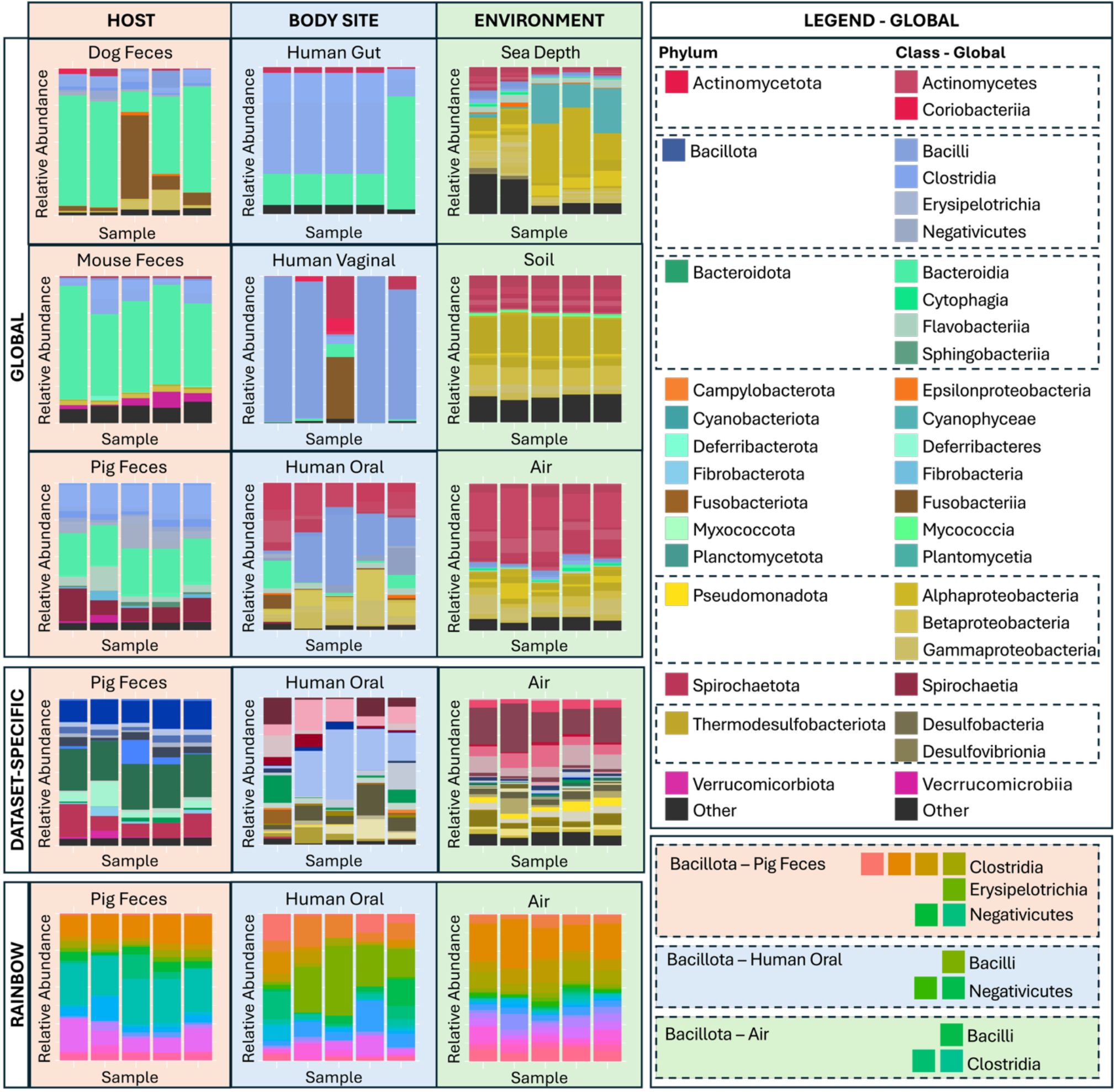
Stacked bar charts indicating relative abundance of taxonomic orders across nine microbiome datasets. For optimal comparison, all taxa were ordered by phylum, class and order membership. Top set of nine panels: PhylaChrome in global mode applied to all nine datasets. Global mode legend includes only phylum-level and class-level colours; all order-level colours are similar to their corresponding class. Middle set of three panels: PhylaChrome in dataset-specific mode applied to three datasets (no legend provided). Bottom set of three panels: Ordered rainbow palette applied to the same three datasets, only the legend for Bacillota orders are provided.

From these charts we see that colours assigned by PhylaChrome global mode allow community structure to be readily compared across multiple taxonomic levels and datasets. For example, at the level of phylum we observe that while Bacteroidota dominate communities in the dog and mouse fecal datasets, samples in the pig dataset contain relatively equivalent abundances of Bacteroidota and Bacillota. Turning to class and order, visualization of the human body sites reveal that the oral dataset contains the largest number of orders within each phylum, but for phyla such as Actinomycetota all orders belong to the same class, as indicated through the saturation and lightness values defined by their parent class; samples from the gut and vaginal datasets show an additional Actinomycetota class but with fewer orders within the phylum. Like the oral dataset, across environmental datasets we observe multiple orders assigned to the same Actinomycetota class.

While global mode allows ready comparisons across datasets, we do note some limitations in visualization, particularly for parent taxa encompassing large numbers of related taxa. For example, the phylum Pseudomonadota contains nine classes resulting in SL-defined colour assignments that can be challenging to differentiate. To overcome this limitation, PhylaChrome dataset-specific mode can be used (**Figure 2**) where taxa within phyla such as Pseudomonadota are more clearly delineated relative to global mode. This is in contrast to the ordered rainbow palette where limited taxonomic information is communicated; in each of the three datasets presented, the colours representing taxa of the same order differed. For example, the Bacillota orders associated with the Pig fecal samples ranged from orange to green, while in the human oral samples, Bacillota orders were associated with only shades of green; ordering taxa by, for example, abundance would further reduce taxonomic information communicated through colour. In all three datasets colour assignments using the rainbow colour palette result in stacked bar charts in which adjacent phyla are challenging to distinguish.

PhylaChrome allows visualization of any bacterial taxa within the NCBI or GTDB database. PhylaChrome can be used to define colours for any graphic where taxa are differentiated through colour. Global mode reflects taxonomic lineage for an entire taxonomic tree allowing intuitive comparisons across datasets, while dataset-specific mode offers the benefit of visually differentiating present taxa within each phylum. PhylaChrome is the first systematic and deterministic colour generator to capture taxonomic relationships for microbiome datasets.

## ONLINE METHODS

PhylaChrome communicates taxonomic information through colour. PhylaChrome uses the (H)ue (S)aturation and (L)ightness (HSL) colour space but provides the user with hex-colour codes. Hue is predefined (either default or user-defined) for phyla, taxa are assigned the hue of their parent phylum and are differentiated by S and L values. Empirically, bounding the allowed S values between 20-100 and the allowed L values between 30-80 produced the most visually distinguishable colours. These are referred to as the global S and L bounds.

### Global Mode

For a given taxonomic database, global mode assigns a deterministic colour to every taxon from phylum level to species level using its scientific name. The colour of a taxon is related to its parent. The S and L values are calculated to differentiate taxa within each phylum. The following definitions are used in the algorithm description:

- 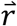 is a vector of requested taxa for which a colour is to be assigned. It is extracted from the user input.
- 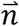 is a vector of all taxa to be assigned a colour, updated programmatically. Initially 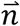 contains all phyla in the lineage of taxa in 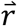. Only phyla for which a hue is assigned are included in 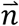.
- 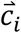 is a vector of all direct children of taxon *i*, exactly one taxonomic level below taxon *i*.
- *h*_*used*_ is a dictionary where the key is a hue and the value is a two-dimensional array of S and L pairings assigned to taxa with the corresponding hue.

The algorithm parses the taxonomic tree using a breadth-first approach; colours are assigned across a taxonomic level before being calculated for child taxa. The algorithm is repeated until either the lowest taxonomic level requested is reached or the species level is reached, whichever is a higher level. Below we illustrate the application of the algorithm to a single phylum called *p* that is an element of 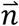.

When *p* is the first entry in 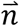, *p* is removed from 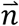 and the hex colour for *p* is extracted from the predefined phylum-level colours. The hex colour code is converted to the HSL colour space using the python package *chromato* (https://github.com/vikpe/chromato). All children of *p* are stored in 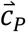 and appended to 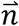. As no taxa within the phylum have been assigned a colour yet, children in 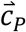 can have a S and L values anywhere in the allowed SL space, defined by the global S and L bounds. This process is repeated for each phylum.

Once the first child of 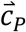, *c*_1_, is at the front of 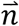, a colour is assigned to *c*_1_ by converting its scientific name to a four-digit number using the SHA-256 cryptographic hash function. A cryptographic has function was selected because an important property of these functions is that two inputs close in value will produce largely different outputs; this is also a desirable property for colour assignment of taxa that may have similar scientific names. The first two digits of the output are used by equation (2) as the value *k* to calculate the S value for *c*_1_. The second two digits are used by equation 2 as the value *k* to calculate the L value of *c*_1_. In equation 2, *v*_*min*_ is the minimum allowed value for S or L, *v*_*max*_ is the maximum allowed value for S or L, and *v*_*final*_ is the final assigned value for S or L. The H value is the same as *p*. The dictionary *h*_*used*_ is updated and the final HSL colour for the *c*_1_ is saved as a new dictionary where the node is used as the key. All direct children of *c*_1_ are appended to the end of 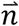. This process is repeated until every child in 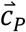 has been assigned a colour.

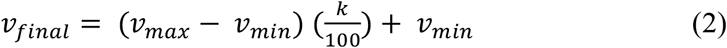

Once the next entry in 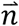 is no longer in 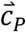, the allowable SL range children of each taxon in 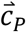 is calculated. From *h*_*used*_ all S and L values used for the hue are extracted. For a given child, *c*_*i*_ the assigned S and L values are *S*_*i*_ and *L*_*i*_. From all currently used S and L values the closest S value that is smaller than *S*_*i*_ and closest S value greater than *S*_*i*_ are extracted as *S*_*less*_ and *S*_*greater*_ respectively. The same is done for *L*_*i*_ to assign *L*_*less*_ and *L*_*greater*_. The minimum allowed S value and L value for children of *c*_*i*_ are calculated using equation (3) and the maximum allowed values are calculated using equation (4). The value is adjusted such that it is within the global S and L ranges and saved for all children of *c*_*i*_. This is repeated for each taxon in 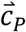.

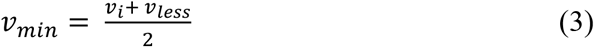

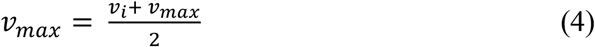

Once the allowable range for children of each taxon in 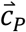 is calculated, 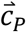 is updated to contain children of the parent phylum for the new first node in 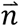. By appending children to only the end of 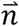 and updating allowable ranges only after all children of the current parent taxon are assigned, the breadth-first approach is implemented.

### Dataset-Specific Mode

Dataset-specific mode assigns colours to all requested taxa of a phylum such that they are maximally separated in the S and L space. The goal of this mode is to optimize the ability to distinguish between assigned colours. Colours are assigned by phylum membership where all taxa of each phylum are assigned at once, regardless of taxonomic rank.

For each phylum, *p*, all requested taxa within the phylum are stored in the vector, 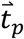 The length of 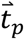 is *m*. From a random uniform distribution, *m* two-dimensional datapoints are selected where one axis is bounded by global S bounds, and the other by global L bounds. A SLSQP algorithm is used to find new coordinates for *m* points that minimize the inverse of the smallest distance between points; a penalty of 10 is added if any two points are within a Euclidean distance of 1 unit. From the final set of *m* points, an inter-point distance matrix *D* is created.

Next, taxa in 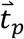 are assigned to the final points from SLSQP. First, the lineage of each taxon in 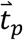 is extracted as a set of taxa. Next, the pairwise Jaccard distance matrix *J* is calculated for all taxa such that entry *J*_*i,j*_ is the Jaccard distance between taxa *t*_*i*_ and *t*_*j*_ with lineage sets *L*_*i*_ and *L*_*j*_ found using equation (5). The python linear sum assignment function is used to perform distance profile matching between *D* and *J*. The cost matrix *C* is calculated such that entry *C*_*i,j*_ is the cost of assigning point *i* to taxon *j*, or the distance between row *i* of *D* and row *j* of *J*. The result is taxa with the most similar lineages are assigned to the closest SLSQP points. The final point a taxon is assigned to is used as S and L values for the taxon.

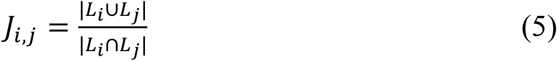

### Additional Functions

Default phylum-level colours were calculated for the NCBI database as of May 2026 and the GTDB database release 226. Only bacterial phyla were considered, and colours were assigned based on phyla associated with the gut microbiome. For different database versions or different microbiomes, the user may consider changing phylum-level colours. To ease this process, PhylaChrome offers three additional functions:

- getPhylaCSV will return a CSV of two columns; the first column contains all phyla within the requested database, and the second column is empty for the user to fill in colours in hex-code format.
- getPhylaCSVBacteria is the same as getPhylaCSV except only phyla in the domain Bacteria are included in the first column.
- getDefaultColours returns the default phylum level colours for the NCBI or GTDB database. The user can then make edits to only some phyla or use the colours for assignments to database versions with different phyla names.

### Dataset Pre-Processing

We applied PhylaChrome to nine shotgun metagenomic datasets from the NCBI. Datasets were selected to compare community structure across hosts, body sites in humans, and environments (**Supplemental Table 1**). For comparison of host datasets, all samples were collected from fecal samples of healthy dogs ^1^, mice ^2^, or pigs ^3^. For comparison of body sites in humans, samples were collected from fecal samples of healthy humans ^4^, oral samples from periodontally healthy humans ^5^, and vaginal samples from women at risk of pre-term birth ^6^. For comparison across environments samples were collected from sea sediments from 5m depth to 750m depth ^7^, air samples from 5 cities ^8^ and soil samples from the same geographical area ^9^. Datasets were processed independently.

For each sample, reads were cleaned, host sequences were removed, and taxonomic classification was performed. Reads were first cleaned and filtered using BBDuk ^10^. For host datasets and body site datasets, host reads were aligned to corresponding NCBI reference genomes (**Supplemental Table 1**) and removed using bowtie2 ^11^. Filtered reads were classified using Kraken2 ^12^ and the standard NCBI database as of April 9^th^ 2026 (https://benlangmead.github.io/aws-indexes/k2). Taxa classifications were collapsed to level order, and abundances were normalized to sum to 1. For each dataset, orders that did not have an abundance of at least 1% in any sample were re-classified as “Other”. For taxon colour assignment, unique orders in each dataset were used as input to PhylaChrome in global mode. For pig feces, human oral, and city air datasets, unique orders were also used as input to PhylaChrome in dataset-specific mode or ordered by taxon membership and mapped to the default rainbow colour scheme in R. Stacked bar charts were created for each dataset using the *ggplot* function in R.

## DATA AVAILABILITY

All source data are available from NCBI at BioProjects and corresponding SRA numbers in Supplemental Table 1. For each dataset, the Kraken2 output is available in the GitHub repository *ParkinsonLab/PhylaChrome*.

## CODE AVAILABILITY

PhylaChrome is available as open-source software under the MIT license in the GitHub repository *ParkinsonLab/PhylaChrome*. The R package can be installed from the GitHub repository. Python code can be downloaded from the GitHub repository and run by importing functions from the script *PhylaChromeMain*.*py*.

## ACKNOWLEDGEMENTS

We thank the members of the Parkinson Lab for extensive discussions of the algorithm during lab meetings. This includes, in no particular order: Ana Popovic, Billy Taj, Gowshigga Thamotharampillai, Irina Utkina, James St Pierre, Grant Stevens, Duncan Carruthers-Lay, Ryan Chieu, Yami Arizmendi Cardenas, Grace Parish, Daniel Fry and Khairatun Yusuff. We would like to further thank Ana Popovic, Yami Arizmendi Cardenas, Grace Parish, and Daniel Fry for testing multiple versions of PhylaChrome in the context of their projects and for providing valuable feedback.

## CONTRIBUTIONS

R.L.T. and J.P. conceived the study and led the study design. R.L.T. developed the tool and performed all the analyses and interpretation. R.L.T. and J.P. wrote the manuscript. J.P. supervised the research. R.L.T. and J.P. read and approved the final manuscript.

