## Supplemental Table 1 for "What colour are Actinomycetes? We have a tool for that! PhylaChrome - A systematic and deterministic tool for assigning colours that capture taxonomic relationships in microbiome datasets"

**Supplementary Material**

**Table S1.** All metagenomic datasets were collected using ILLUMINA sequencers in paired mode. SRA numbers are listed for the five samples used from each dataset, in order of appearance from left to right in the corresponding chart in **Figure 2.**

| **Dataset** | **BioProject** | **NCBI Host Reference Genome** | **Sequencer** | **SRA Numbers** | **Description** |
|---|---|---|---|---|---|
| Host | | | | | |
| Dog | PRJNA1222153 | GCF_011100685.1 | HiSeq 2500 | SRR32305984<br>SRR32305991<br>SRR32306002<br>SRR32306013<br>SRR32306018 | All healthy controls |
| Mouse | PRJEB18589 | GCF_000001635.27 | HiSeq 2500 | SRR12506358<br>SRR12506359<br>SRR12506360<br>SRR12506361<br>SRR12506362 | All healthy controls |
| Pig | PRJNA629856 | GCF_000003025.6 | NovaSeq 6000 | SRR11808123<br>SRR11808126<br>SRR11808128<br>SRR11808129<br>SRR11808130 | All healthy controls |
| Body Site | | | | | |
| Human Gut | PRJEB6070 | GCF_000001405.40 | NextSeq 550 | ERR478964<br>ERR478965<br>ERR478966<br>ERR478967<br>ERR479570 | All age 34-35, diagnosis “normal” |

| | | | | | |
|---|---|---|---|---|---|
| Human Oral | PRJNA1183294 | GCF_000001405.40 | NovaSeq 6000 | SRR31350376<br>SRR31350377<br>SRR31350388<br>SRR31350399<br>SRR31350410 | All diagnosed periodontally healthy |
| Human Vaginal | PRJEB34536 | GCF_000001405.40 | NextSeq 500 | ERR4421550<br>ERR4421551<br>ERR4421552<br>ERR4421553<br>ERR4421554 | All pregnant women at risk of pre-term birth. Metadata not available. |
| Environment | | | | | |
| Sea Sediment | PRJNA638805 | N/A | HiSeq X Ten | SRR12042682<br>SRR12042683<br>SRR12042684<br>SRR12042685<br>SRR12042686 | 750 m depth<br>150m depth<br>30 m depth<br>5m depth<br>5 m depth |
| Soil | PRJNA779554 | N/A | NovaSeq 6000 | SRR16915451<br>SRR16915453<br>SRR16915458<br>SRR16915465<br>SRR16915470 | All first collection time point |
| City Air | PRJNA561080 | N/A | HiSeq 2500 | SRR10002674<br>SRR10002678<br>SRR10002689<br>SRR10002695 | Sweden<br>Hong Kong<br>USA<br>Norway<br>UK |

| | | | | |
|---|---|---|---|---|
| | | | | SRR1000269 9 |
